# U-FISH: a universal deep learning approach for accurate FISH spot detection across diverse datasets

**DOI:** 10.1101/2024.03.06.583706

**Authors:** Weize Xu, Huaiyuan Cai, Qian Zhang, Florian Mueller, Wei Ouyang, Gang Cao

## Abstract

In the rapidly advancing landscape of fluorescence in situ hybridization (FISH) technologies, there is a critical need for sophisticated yet adaptable methods for spot detection. This study introduces U-FISH, a deep learning approach that significantly improves accuracy and generalization capabilities. Our method utilizes a U-Net model to transform noisy and ambiguous FISH images into a standardized representation with consistent signal characteristics, facilitating efficient spot detection. For the training and evaluation of the U-FISH model, we have constructed a comprehensive dataset comprising over 4,000 images and more than 1.6 million manually annotated spots, sourced from both experimental and simulated environments. Our benchmarks demonstrate that U-FISH outperforms existing methods for FISH spot detection, offering improved versatility by eliminating the need for laborious manual parameter adjustments. This allows for its application across a broad spectrum of datasets and formats. Furthermore, U-FISH is designed for high scalability and is capable of processing 3D data, supporting the latest generation of file formats for large and complex datasets. To promote community adoption and ensure accessibility, we provide a user-friendly interfaces: Napari plugin, web application and command-line interface. The complete training dataset is made publicly available, laying a solid foundation for future research in this field.

## Introduction

Imaging technologies play a pivotal role in biomedical research. In the study of gene regulation, methods with single-molecule sensitivity have provided invaluable insight to our understanding of how individual cells regulate their genome in space and time [1]. Fluorescent In Situ Hybridization (FISH) techniques are instrumental approaches permitting the visualization of individual RNAs or genomic loci with high spatial resolution in their native cellular environment. Initial implementations of single-molecule FISH (smFISH) are used to probe a handful of different mRNA species [2, 3]. Recent advancements in probe synthesis, multiplexed barcoded detections, signal amplification and tissue clearing, opened the door for the field of spatial transcriptomics and spatial genomics. These approaches now enable the visualization of hundreds to thousands of genes with single-molecule sensitivity in complex tissues at 3D level[4, 5, 6, 7, 8]. However, the analysis of this increasingly large and complex datasets remains a challenge, in particular, the accurate and efficient detection of diffraction-limited RNA spots, which is especially challenging in images with varying background levels and spot intensities. Many rule-based methods have been developed to address the challenge of spot detection in microscope images[9, 10, 11, 12, 13]. However, a significant limitation of these methods lies in their inability to uniformly apply across different datasets without parameter adjustments. This limitation stems from the varying imaging conditions and sample characteristics across datasets, which introduce distinct features requiring time-consuming parameter tuning for each image.

In the evolving field of biomedical imaging, deep learning has revolutionized both traditional methods and introduced novel approaches for image analysis. Technologies such as Cellpose[14, 15] for cell segmentation, CARE[16, 17] for image reconstruction, and advanced techniques for cell image classification[18] stand as pillars of this transformation. Each of these applications demonstrates deep learning’s capacity to not only simplify but also significantly enhance the precision and efficiency of analyzing microscopic images. Building upon these foundational achievements, deep learning has further extended its influence to the specific challenges of spot detection in microscopy images. This advancement is crucial for understanding cellular behaviors and gene expression patterns at a granular level. Techniques such as DetNet[19], SpotLearn[20], and deepBlink[21] leverage deep learning models to achieve FISH spot detection without the need for manual parameter tuning. This automation represents a significant advancement over traditional methods, streamlining the detection process and enhancing accuracy. Additionally, DeepSpot[22] specializes in the enhancement of RNA spots in single-molecule FISH (smFISH) microscopy images. However, when it comes to applying these methods in real-world scenarios, there are notable shortcomings that remain to be addressed. First and foremost, there is a lack of sufficiently broad datasets, which hampers the training of versatile models capable of effectively dealing with FISH data from diverse sources. Additionally, these methods still exhibit gaps in their ability to process 3D data and in terms of usability, making the transition from theoretical application to practical use somewhat challenging.

Here, we present U-FISH, an advanced FISH spot detection tool based on deep learning. We designed U-FISH to overcome existing challenges in FISH image analysis and establish a universally applicable algorithm for rapid and precise spot detection, adapted to a multitude of FISH imaging modalities. To achieve this objective, we first created a dataset comprising over 4,000 images, totaling more than 1.6 million annotated spots, obtained from diverse sources, including both experimental and simulated data(Fig.1 and Method Dataset construction). We then developed an optimized signal enhancement network designed to augment input images of varying characteristics(Fig.2, Extended Data Fig.1 and Method U-FISH model design). The enhanced images produced by this network exhibit high signal-to-noise ratios for smFISH spots and possess consistently stable features, facilitating more accurate spot detection. It’s eliminated the need for laborious manual parameter tuning across datasets. When compare with existing deep-learning solutions[20, 19, 21], U-FISH not only achieves a higher accuracy but also boasts a smaller model size and faster inference(Fig.3 and Extended Data Fig.2). Furthermore, U-FISH is specifically designed to accommodate 3D data and is compatible with large data storage formats(OME-Zarr[23] and N5), thus providing a unified, scalable solution for smFISH image analysis(Fig.4). To facilitate the widespread adoption of U-FISH within the scientific community, we have developed the U-FISH application(Fig.5). In this application, a range of accessible interfaces are included: Napari[24] plugin, web application(ImJoy[25] plugin, available at https://ufish-team.github.io/), command-line interface (CLI) and API(https://github.com/UFISH-Team/U-FISH). Lastly, to foster open science and encourage the development of related tools, the comprehensive dataset utilized in this study is publicly accessible at https://huggingface.co/datasets/GangCaoLab/FISH_spots, aiming to support future advancements and enhance reproducibility.

**Fig.1:**
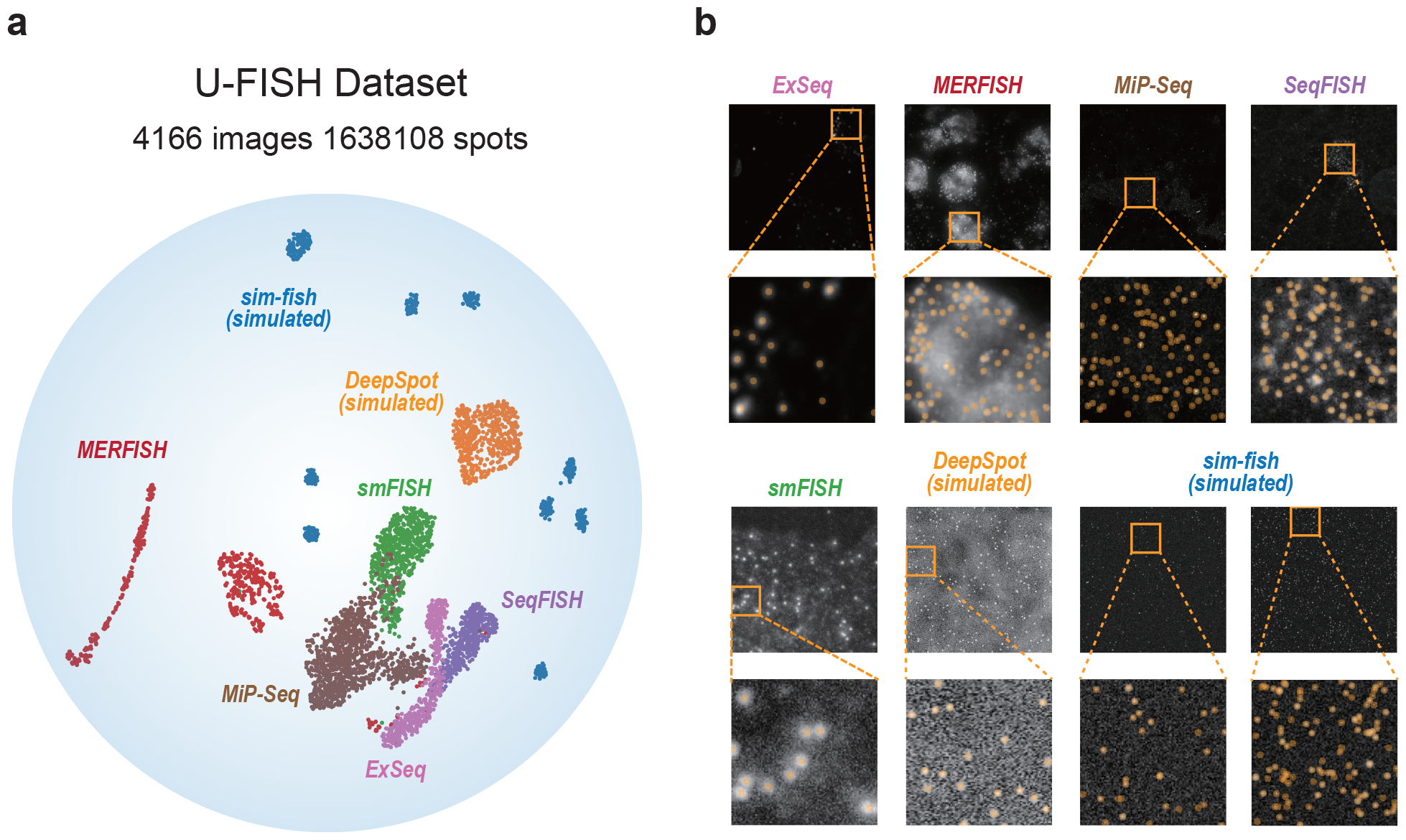
U-FISH dataset. **a**, scatter plot illustrating the UMAP dimensionality reduction of the images in the U-FISH dataset. This diverse dataset encompasses a total of 4,166 images sourced from seven different origins, featuring 1,638,108 manually verified spots. UMAP visualization captures the dataset’s inherent diversity, showcasing the variance in data distribution and complexity across different sources. **b**, Sample images from each of the seven distinct sources, accompanied by annotations of spot coordinates, are displayed. These examples highlight the variability in background characteristics and signal spot features among the different sources. This diversity underscores the comprehensive nature of the U-FISH dataset, ensuring its robustness and applicability across a wide range of FISH imaging scenarios.

**Fig.2:**
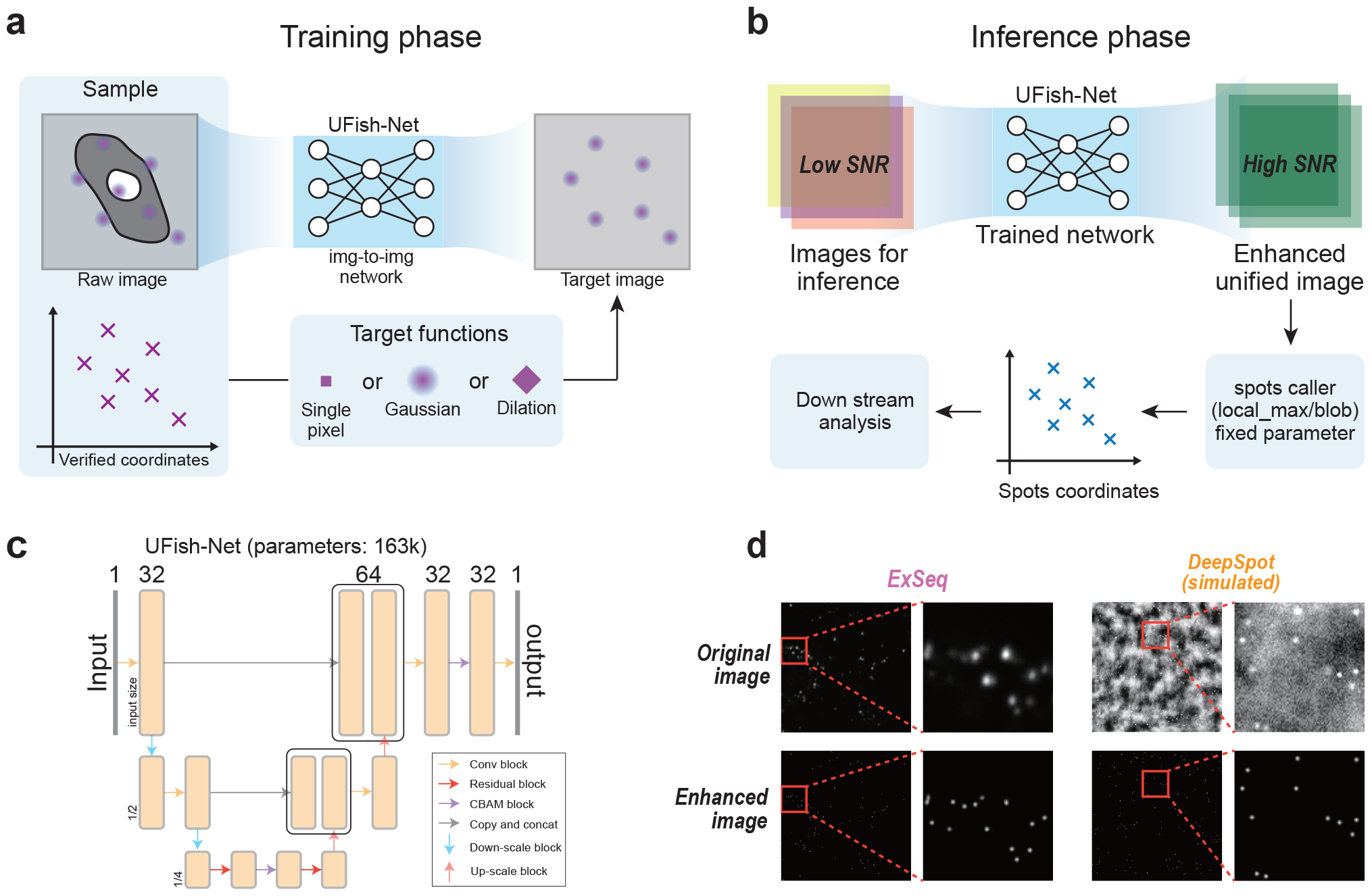
U-FISH enhances the original images into a standardized representation with consistent signal characteristics for efficient spot detection. **a**, During training, the model leverages coordinate information from annotations to generate the target images. At this stage, we provide configurable options for the type of target function to be applied, including single pixel, Gaussian, and dilation. However, for the analyses presented in this paper, we primarily utilize the Gaussian option. The img-to-img model is then trained using these target images alongside the original images to learn the transformation. **b**, During the inference phase, the trained model inputs original images, typically characterized by low signal-to-noise ratio (SNR), and outputs enhanced images with consistent features and significantly reduced background noise, resulting in a higher SNR. Spot detection is then executed on these enhanced images using a fixed-parameter, rule-based algorithm (see Methods Spot detection on enhanced images), which is selected based on the target function used during training. This process efficiently identifies spot coordinates. **c**, UFish-Net architecture is based on a U-Net with depth two, featuring two rounds each of downsampling and upsampling, and the integration of CBAM(Convolutional Block Attention Module) at the bottom and in the final decoder layers. This network is notably compact, with only 163k parameters. **d**, Comparative visualization between original and enhanced images demonstrates the increased SNR in the enhanced images. The enhancement process effectively transforms signal points with varying disticnt characteristics into uniformly featured spots, as shown in the enhanced images.

**Fig.3:**
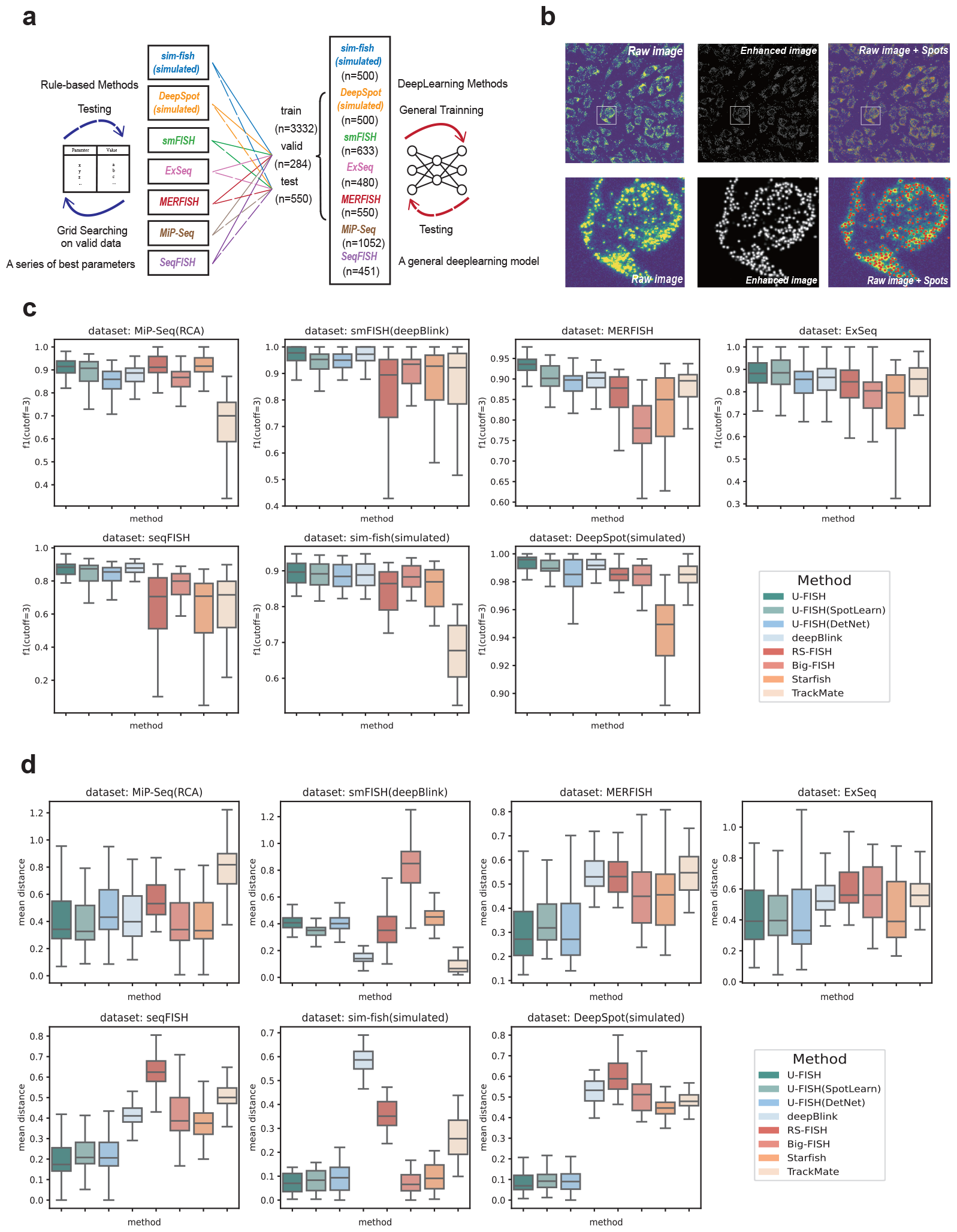
Benchmark of different spot detection software. **a**, An illustrative diagram delineates the benchmarking strategy employed. For rule-based methods, optimal parameters are identified through a grid search conducted on the validation dataset. These optimized parameters are then applied to the test dataset to evaluate performance. In contrast, deep learning-based models are trained on the training dataset, with the validation dataset used for model selection before performance testing on the test dataset. **b**, example showcasing U-FISH’s capability in image enhancement and detected signal spots. This illustration demonstrates the effectiveness of U-FISH in enhancing image quality and accuracy in spot identification. **c**, box plots displaying the performance of various models across datasets from different sources, measured by the F1 score (cutoff=3). A higher F1 score indicates a stronger accuracy in spot detection. The U-FISH (DetNet) and U-FISH (SpotLearn) in legend represent the implementation of these two networks in the U-FISH framework, as described in Benchmark for deep learning methods. **d**, box plots as in c, but focusing on the mean distance metric. This metric represents the average distance between the coordinates of detected signal spots and their nearest true signal points. A lower mean distance implies a smaller localization error, showcasing the precision of the spot detection method. These comparative analyses provide a comprehensive overview of U-FISH’s performance against other spot detection methodologies, highlighting its proficiency in accuracy.

**Fig.4:**
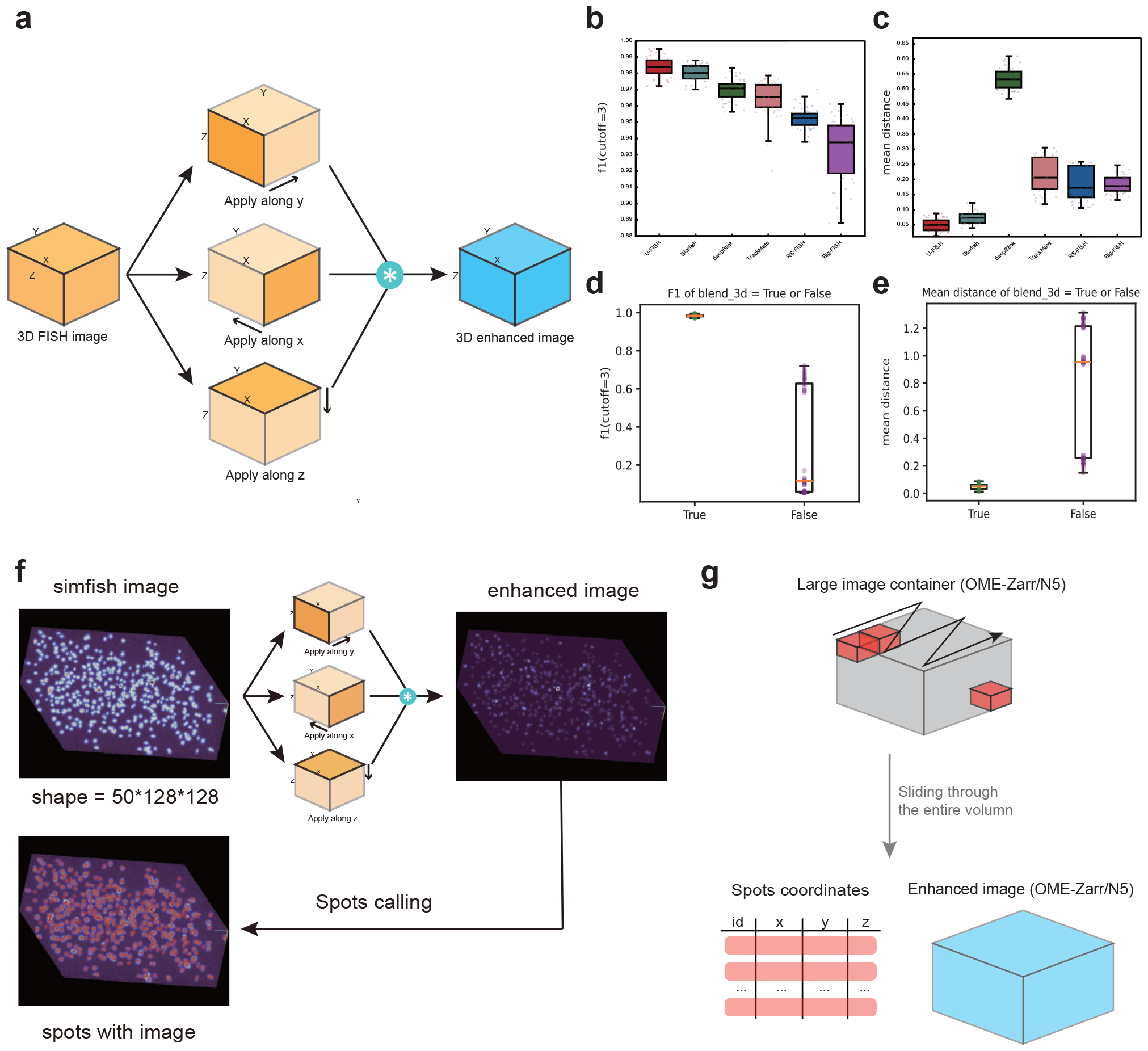
3D Inference Strategies and Benchmarks in U-FISH. **a**, schematic representation of the 3D blending strategy used for volumetric images. In this approach, the U-FISH model is applied along the three axes (x, y, z) for image enhancement. The resultant images from each axis are then multiplied together to reconstruct a 3D enhanced image. **b**,**c** The performance of different methods on a 3D benchmark using simulated datasets. **b**, The differences in F1 scores. **c**, The variations in mean distances among the methods. **d** and **e** the accuracy of the results obtained using the 3D blending strategy against enhancement performed solely along the z-axis. Here, **d** focuses on the F1 score metric, and **e** on the mean distance metric. **f**, Example of 3D data enhancement and spot detection using the 3D blend strategy, showcasing its practical application. **g**, Strategy employed by U-FISH for large-scale image inference. Typically, large images are stored in formats like OME-Zarr on a disk. U-FISH efficiently handles these by reading and processing individual data blocks at a time, storing the results back to the disk, and then sliding the window to process the entire dataset. This method ensures efficient handling and analysis of large-scale datasets, demonstrating the adaptability and robustness of U-FISH in various imaging scenarios.

**Fig.5:**
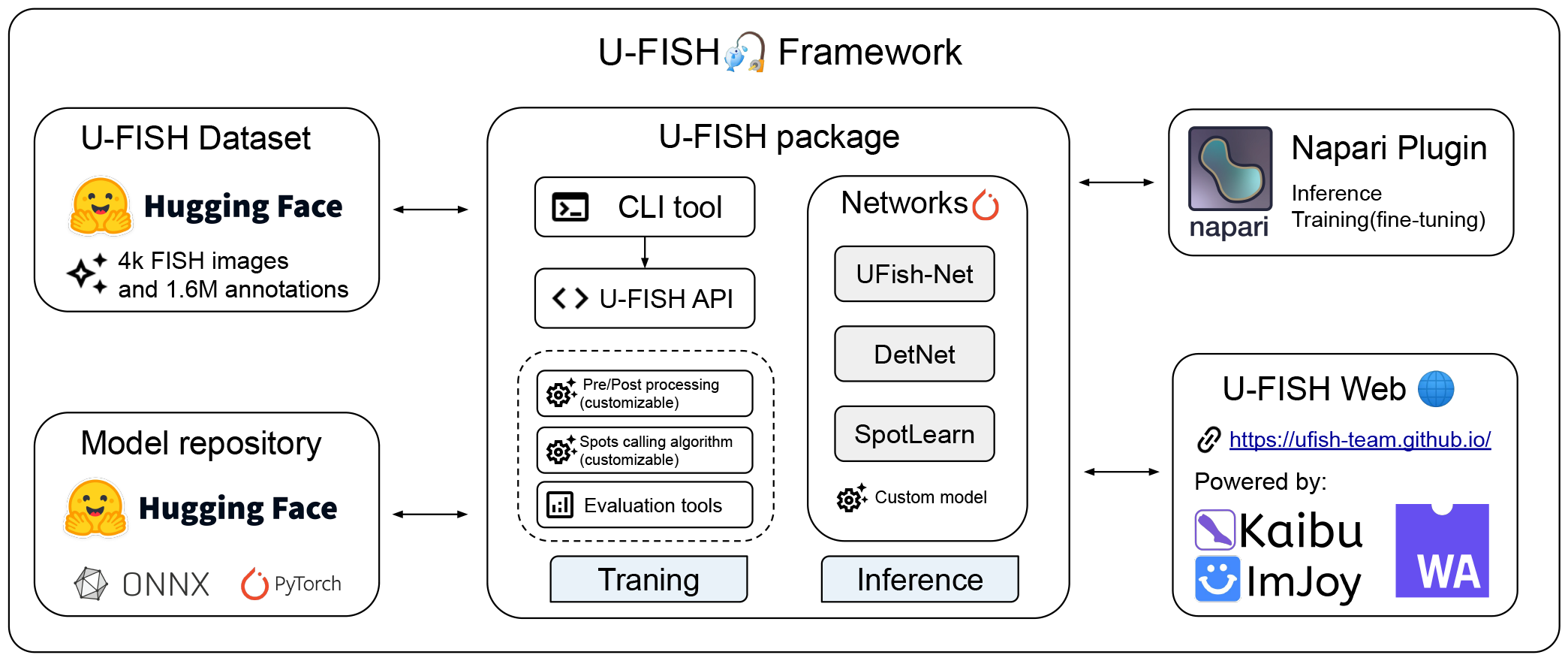
The U-FISH Framework provides a comprehensive foundation for the development of future FISH spot identification methods, comprising several key components. (1) U-FISH Dataset: A publicly available dataset hosted on HuggingFace, featuring a vast collection of raw images and annotated FISH spots data ready for immediate use. (2) Model Repository: A collection of network weights that have been pre-trained on the U-FISH dataset is also publicly accessible on the HuggingFace platform. These weights can be directly utilized by users or serve as a basis for model fine-tuning. (3) U-FISH Package: This Python package includes the U-FISH API, CLI, and network architecture definitions (with support for custom network architectures by users). It encompasses a suite of modular components for image pre-processing, post-processing, spot detection algorithms, and result evaluation tools. (4) Napari Plugin: It enables users to invoke U-FISH for predictions and fine-tuning directly within the Napari viewer using a graphical user interface. (5) U-FISH Web: A web application that allows users to use U-FISH for predictions without the need to install any dependencies, accessible directly through a web browser. Together, these components make the U-FISH Framework a versatile and user-friendly toolkit for researchers and practitioners in the field of FISH analysis.

## Result

### U-FISH enabling universal, rapid, and precise detection of FISH spots

To ensures consistent detection performance across different FISH datasets without the need for manual parameter adjustments, U-FISH employs a U-Net model[26] to transform raw smFISH images with varying characteristics into enhanced images with uniform signal spot characteristics and an improved signal-to-noise ratio, on which it then detects signal spots without time-consuming manual parameter tuning(Fig.2a,b and Methods). One of U-FISH’s cornerstone features is its comprehensive training dataset, comprised of over 4000 images with more than 1.6 million verified targets from seven diverse sources. This data diversity allows to train a universal spot detection model and for an unbiased evaluation of the algorithm’s performance (Fig.1). Detailed information on the dataset and model specifics are provided in the Methods section.

To rigorously test U-FISH’s performance, we conducted a detailed comparison across various datasets. To benchmark its spot detection efficiency, we used both experimental and simulated data. For quantitative performance comparison, we used F1 scores and mean distance errors on the test dataset and comparing these metrics against several other methodologies, including both deep learning-based and rule-based approaches (Fig.3a and Method Benchmark for deep learning methods, Benchmark for rule-based methods). Remarkably, U-FISH demonstrated exceptional performance across these diverse datasets without the necessity for dataset-specific parameter tuning, showcasing its robustness and versatility. In fact, U-FISH achieved the best performance on the majority of datasets compared to the evaluated methods. For the mean metric scores, U-FISH achieved an F1 score of approximately 0.908, surpassing deepBlink (F1: 0.890), DetNet (F1: 0.871), SpotLearn (F1: 0.890), Big-FISH (F1: 0.830), RS-FISH (F1: 0.841), Starfish (F1: 0.829), and TrackMate (F1: 0.765).

In terms of mean distance, U-FISH recorded a value of 0.311, outperforming all competitors except for DetNet (mean distance: 0.334) and SpotLearn (mean distance: 0.305), indicating its high precision in spot localization (For more detailed benchmarks on all datasets shown in Fig.3c,d). Furthermore, our tests on datasets with simulated noise revealed that U-FISH possesses strong noise resistance capabilities (Extended Data Fig.2f,g), highlighting its suitability for analyzing FISH images under various experimental conditions and diverse samples from diverse biological origins.

Generalization capability is a cornerstone of deep learning models, crucial for their effectiveness and applicability across various domains. It enables models to perform well not only on the data they were trained on but also on new, unseen datasets. To assess the generalization capability of models across diverse types of datasets, we compared performance of three distinct training modalities: General (trained on all datasets), Special (trained only on one specific dataset), and Fine-Tuned (retrained General model for a specific dataset), see Extended Data Fig3a. The outcomes of this comparative study reveal that, overall, the performance gaps among the three models are relatively small. However, in certain datasets, the General and Finetune models demonstrated superior performance over the Special model (Extended Data Fig.3b). This suggests that training on a diverse array of datasets enables the model to achieve enhanced versatility, likely due to exposure to a wider variety of features and patterns in different kinds of source datasets. Additionally, we conducted a leave-one-out experiment to test the model’s generalization ability on unseen datasets. In this experimental setup, data from one source was excluded from the training process and then used as a test set to evaluate the model’s performance(Extended Data Fig.3c). The results of this test indicate that the U-FISH model(mean F1: 0.834, mean distance: 0.429) exhibits superior generalization capabilities compared to deepBlink(mean F1: 0.823, mean distance: 0.531), with better performance in adapting to and accurately processing data which had not encountered during training(Extended Data Fig.3d). This further underscores U-FISH’s robustness and its potential as a versatile tool for FISH image analysis across a wide range of dataset characteristics.

To further explore the data requirements for training a General model from scratch and the impact of training data volume on performance, we conducted downsampling tests on the number of training images. The results indicate that a satisfactory performance (average F1 score *>* 0.85) can be achieved with approximately 300 training images(with 512×512 pixel size) when retraining the model from scratch. Moreover, when the number of training images exceeds 1800, the model performance further improves (average F1 score *>* 0.9), with the F1 score showing a slow upward trend with the dataset size increases thereafter(Extended Data Fig.3e). Curating such a larger number of annotated images can be daunting task. We therefore wanted to investigate if parameter fine-tuning based on the pre-trained weights of our General model could reduce the size of the required training data. For this purpose, we wanted to use a challenging dataset, and turned to images of single RNA translation with the SunTag approach [27]. Importantly, these images also feature distinct spots, but the experimental modality is different from smFISH. Our findings reveal that fine-tuning on the General model requires fewer training images compared to training a model from scratch to achieve better performance. Remarkably, approximately only about 30 training images (with a resolution of 512 × 512 pixels) were sufficient to achieve commendable results through fine-tuning, and superior accuracy was maintained even with increased number of training images (Extended Data Fig.3f).

In summary, our findings underscore the value of leveraging training strategies on diverse data-sources to boost model performance across a spectrum of different datasets, highlighting the potential benefits of a more universal approach to model training in achieving robust generalization capabilities. Lastly, it underscores the practical advantages of fine-tuning from a well-established General model base.

Lastly, U-FISH’s optimized and compact network architecture, comprising only 163k parameters, contributes significantly to its computational efficiency. This streamlined design not only ensures that U-FISH maintains exceptional accuracy but also achieves superior processing speeds, particularly on devices equipped with GPUs, making it an ideal choice for high-throughput analysis. Its commendable efficiency extends to CPU-based systems as well (Extended Data Fig.2a,b), demonstrating the model’s versatility across various computing environments. The compact size of the U-FISH network, with an ONNX file merely 676kB in size, underscores the ease with which it can be deployed, even in resource-constrained settings. This level of computational performance and small footprint underpin U-FISH’s capability to effectively handle large images and broadens its applicability across a variety of scenarios, including deployment in web browsers. The combination of high efficiency, low parameter count, and minimal storage requirements makes U-FISH a highly deployable solution, suitable for a wide range of applications from desktop-based analysis to cloud and web-based platforms.

### Extends U-FISH to 3D FISH signal detection

An advancement presented by U-FISH is the extension of deep learning-based spot detection methods to 3D imaging, which had not been well resolved before due to certain inherent challenges. One obstacle to applying deep learning approaches to 3D FISH images for spot detection is the difficulty in obtaining high-quality annotated 3D datasets. This challenge is largely attributed to the labor-intensive process of manually identifying center points within the 3D space. The workload required for accurate annotation in 3D is significantly greater than that for 2D images, complicating the acquisition of reliable 3D training data. Additionally, there is a need for efficient implementations that minimize GPU memory requirements for model training, further complicating the transition to 3D. These challenges underscore the necessity of developing a strategy that enables models trained on 2D data to be effectively applied to 3D imaging scenarios, thereby overcoming the hurdles of data annotation and computational demands.

A straightforward approach to applying a 2D model to 3D data involves sliding the model along the z-axis to perform inference. However, since these models are not trained specifically on 3D data, they might inadvertently enhance signals that are not central on the z-axis. This issue can lead to the elongation of signals along the z-axis, potentially causing repetitive detection of a single point when applying downstream detection algorithms. To address this issue, we implemented a new approach, which involves applying 2D models along the three axes (x, y, and z) to enhance the image, resulting in three separate image stacks. These stacks are then multiplied together to reconstruct a 3D enhanced image(Fig.4a,f). Experimental results have demonstrated that this method significantly improves detection accuracy on 3D simulated datasets (Fig.4d,e). Furthermore, comparative analysis with specific benchmark numbers indicates that U-FISH significantly outperforms other methods in terms of accuracy when applied to simulated datasets. For the mean metric scores, U-FISH achieved an F1 score (cutoff=3) of 0.984 and a mean distance of 0.049, demonstrating superior performance in spot detection accuracy and precision of localization. In comparison, deepBlink recorded an F1 score of 0.970 and a mean distance of 0.533, while Big-FISH showed an F1 score of 0.933 and a mean distance of 0.182. RS-FISH and TrackMate also trailed behind with F1 scores of 0.952 and 0.964, and mean distances of 0.186 and 0.213, respectively. Notably, Starfish came close with an F1 score of 0.980 and a mean distance of 0.072. These results, illustrated in Fig.4b and c, underscore U-FISH’s advanced capability in detecting FISH signals with remarkable accuracy and minimal localization error, setting a new benchmark for performance in simulated 3D datasets.

Building on the innovative approach for 3D spot detection, U-FISH also employs a strategic method for handling large-scale multidimensional-image inference, particularly crucial for large images stored in formats such as OME-Zarr[23]. This strategy involves reading and processing individual data blocks from the disk, one at a time, and then storing the processed results back onto the disk(Fig.4g). The process continues by sliding the window to sequentially handle the entire dataset. This method minimizes memory usage, making U-FISH highly adaptable and robust across various imaging scenarios. This approach further demonstrates U-FISH’s capability to seamlessly integrate into existing bioimaging workflows, providing a scalable solution for the analysis of extensive FISH imaging data, thereby reinforcing the framework’s utility in advancing the field of FISH analysis.

### User-friendly tools for facilitate the community adoption

Next, we extended U-FISH from a model to a comprehensive framework for training of FISH spot detection models, evaluating results, and facilitating inference across multiple platforms (Fig.5). To enhance accessibility for end-users, we developed a user-friendly graphical user interface (GUI) for U-FISH, aiming to streamline the user experience for both novices and experienced researchers. Our Napari Plugin enabling users to invoke U-FISH for predictions and model training directly through a desktop application (Extended Data Fig.5a). In addition to the desktop application, we implemented a U-FISH Web interface, leveraging ImJoy[25], Kaibu, and Webassembly. This innovative approach allows for model predictions to be executed directly within a web browser, significantly lowering the barrier to entry for all users(Extended Data Fig.5.b). The web interface not only makes easy access to U-FISH’s powerful capabilities but also incorporates a visualization of the training data. More specifically, it permits simultaneous display of FISH images and their UMAP dimensionality reductions providing an interactive interaction with these data (Extended Data Fig.1.b).

The core components of U-FISH are encapsulated within a Python package, which offers APIs and CLI interfaces for training, prediction, and result assessment. This package includes network architectures for UFish-Net (Our network), DetNet, and SpotLearn, all implemented using PyTorch, and allows for the integration of custom network architectures by users. Importantly, U-FISH supports a wide range of file formats, including TIFF, OME-Zarr[23], and N5, making it versatile for input data types. It is also capable of handling multi-dimensional images, from 2D (x, y) to 4D (tczxy), ensuring comprehensive applicability across various imaging scenarios. To accommodate large-scale imaging datasets, U-FISH is equipped to process large images through block processing, enabling the handling of images that exceed the available memory size. This approach allows for efficient and scalable analysis, even in resource-constrained environments. The U-FISH package facilitates the easy substitution and modification of functions related to image pre-processing and the post-processing of enhanced images via its API. Its modular design not only enhances the reusability of the U-FISH framework but also greatly benefits subsequent research by facilitating methodological improvements and accelerating the deployment process. Hosted on GitHub with extensive documentation and test data, the Python package can be easily installed via pip. Through these interface solutions, U-FISH aims to democratize access to advanced FISH signal detection and analysis, making it more accessible and user-friendly for the scientific community. This inclusivity ensures that U-FISH stands as a formidable tool in the arsenal of researchers seeking to push the boundaries of FISH analysis.

## Discussion

We introduced U-FISH, a deep learning-based framework designed to meet the vast growing demands for sophisticated spot detection in next generation fluorescence in situ hybridization (FISH) technologies in the spatial-omics era. By leveraging a comprehensive dataset with over 4,000 images and more than 1.6 million manually annotated spots, U-FISH showcases improvement in accuracy and generalization ability. This dataset, encompassing both experiments and simulations, allows U-FISH to operate with high versatility across various datasets and formats without the need for manual parameter adjustments. Furthermore, its design emphasizes scalability, accommodating large and complex data storage formats and extending its utility to 3D FISH data analysis. To improve community adoption and ensure accessibility, U-FISH is complemented by a suite of user-friendly tools, including a Napari plugin, a web application, a command-line tool and the API. The entire dataset is made publicly available, laying a solid foundation for ongoing and future research in this field.

Despite its significant improvements, U-FISH has several limitations that need further attention. One notable challenge lies in the identification of densely packed signal spots, where the current model may struggle to distinguish closely situated points accurately. This limitation underscores the need for further refinement in the algorithm’s ability to resolve high-density areas without sacrificing overall performance. Additionally, while we have built a relatively broad dataset, it may still fall short of covering all scenarios encountered in the real world, leading to sub-optimal performance under certain conditions. Moving forward, expanding the dataset and developing tools for real-time annotation and training could lower the barriers to model fine-tuning, addressing these limitations. Moreover, compared to rule-based methods, deep learning approaches, including U-FISH, are susceptible to the “hallucination” problem, where neural networks may produce false positives in certain situations. This issue, coupled with the inherent “black box” nature of neural networks, makes it challenging to address these false positives through parameter adjustments alone. Overcoming this drawback requires a multifaceted approach, including further refining the model’s architecture, enhancing training methodologies, and possibly integrating interpretability features to demystify the decision-making process of the neural network. Addressing these limitations will be crucial for elevating U-FISH’s accuracy and reliability in FISH signal detection and analysis, ensuring its broader applicability and effectiveness in real-world scenarios.

In summary, U-FISH represents a significant step forward in FISH analysis, offering an adaptable, accurate, and user-friendly solution for researchers. While it pushes the boundaries of what is currently possible in spot detection algorithms, ongoing efforts to address its limitations will be crucial in fully realizing its potential and expanding its applicability to even more challenging datasets. Future enhancements to U-FISH will focus on improving the detection of densely packed spots, extending the framework to incorporate more advanced deep learning models, and enlarging the training dataset to cover a broader range of FISH technologies and experimental conditions. Additionally, efforts will be made to streamline U-FISH’s integration with existing bioinformatics workflows, enabling seamless analysis of FISH data in conjunction with other genomic and transcriptomic datasets. Through these improvements, U-FISH aims to remain at the forefront of innovations in FISH analysis, supporting the scientific community in unraveling the complexities of cellular function and gene expression patterns.

## Methods

### U-FISH model design

Our design draws inspiration from the established image-to-image model architectures, with U-Net[26] being the most extensively utilized among them. This model is renowned for its effectiveness in a diverse array of tasks where image-to-image translation is paramount, providing an excellent foundation for our modified approach.

Following this established protocol, the U-FISH network architecture(UFish-Net) introduces several strategic adjustments to the conventional U-Net model to suit the specific requirements of fluorescence in situ hybridization (FISH) spot detection. By paring down the number of convolutional channels to a constant 32 across all intermediary layers, the architecture achieves a reduction in the complexity and an increment in the speed of computations without compromising on performance.

During the downsampling phase, our network replaces the MaxPooling operation with a strided convolution of 3×3 kernels, resulting in a more refined reduction of spatial dimensions. For upsampling, the choice of nearest neighbor interpolation over transposed convolution has been made to avoid common upsampling issues like checkerboard artifacts. To further enhance the model’s predictive strength, residual connections are introduced post-downsampling and within the base layer of the decoder, ensuring effective feature propagation. Additionally, by incorporating the Convolutional Block Attention Module (CBAM) [28] at strategic decoding junctions, the UFish-Net emphasizes relevant feature sets through attention-focused processing, boosting its accuracy.

### Dataset construction

This dataset is a comprehensive collection of 4,166 fluorescence in situ hybridization (FISH) images, amalgamating real experimental data from a variety of sources including MERFISH, seqFISH, ExSeq, Mip-Seq and smFISH(Fig.1) It also encompasses simulated data from sim-fish and DeepSpot[22] simulators.

The initial phase of the process involved training a standard U-Net convolutional neural network (CNN) using the PyTorch framework. Although this model, with its approximately 8 million parameters, was not intended for practical deployment due to its size, it effectively captured the intricate features of FISH signals and served as a pre-labeling tool for the entire dataset. To ensure the highest level of accuracy and reliability, each image, along with its preliminary annotations generated by the U-Net model, was examined and verified by human annotators. This was conducted using the Napari visualization tool[24], facilitating a rigorous review process where automated model outputs were closely aligned with actual ground truths. This comprehensive manual verification and data curation process not only guaranteed the precision of the annotations but also allowed for the organization and association of metadata with each image, providing detailed insights into experimental conditions and source information(Extended Data Fig.1a).

The final U-FISH dataset, a substantial assembly of 4,166 images, features a total of 1,638,108 annotations. This rich compilation covers an extensive range of experimental conditions and hybridization targets, making it a valuable resource for the development of a generalized FISH point detection algorithm.

### Spot detection on enhanced images

The potency of deep learning models lies in their ability to transform images from varied sources into a representation with consistent features, herein referred to as enhanced images. Given this uniformity, the task of spot detection on enhanced images permits the employment of relatively straightforward and parameter-fixed methods. The choice of spot detection technique is contingent upon the method used for generating target images during the training of the model.

For models trained with single-pixel and Gaussian-smoothed target images, we identify local maxima on the enhanced images using the `skimage.morphology.local_maxima` function. Subsequent to this, a fixed threshold(default 0.5) is applied based on the intensity values at the identified coordinates to sieve through the potential spots, filtering out those below the threshold.

Conversely, for models trained on target images processed with dilation, we first convert the enhanced images into binary images using a set threshold(default 0.5). We then detect all connected components within the binary image. Connected components exceeding a certain size threshold—determined by the area of the structuring element used during dilation—are further segmented using the watershed algorithm. For each identified connected component, the centroid is calculated and designated as the spot’s coordinates.

This methodology affords a streamlined approach to spot detection, circumventing the need for extensive parameter tuning and enabling consistent performance across datasets derived from disparate imaging conditions.

### Training sample construction and network training

For the structuring of our training database, we allocated the images into distinct sets: 3,332 for training, 284 for validation, and 550 for testing. Each data entry is comprised of a pair: an image file in TIFF format and a corresponding spot coordinate file in CSV format. During training, these pairs are read concurrently, with the spot coordinates utilized to produce the target image.

Our experimentation encompassed three distinct methodologies to create these target images: single-pixel representation, dilation, and Gaussian smoothing. The single-pixel target images were crafted by initializing a blank image with identical dimensions to the input image and assigning a value of one to pixels at the spot coordinates. Building upon this, we expanded the marked spots using the `skimage.morphology.dilation` function, with `skimage.morphology.disk` as the structuring element. We tested various sizes of the target to optimize the dilation process. For Gaussian smoothing, the single-pixel image underwent a filtering process with a Gaussian filter set to a sigma value of 1. Post-smoothing, the resulting image’s intensity was normalized to 0-1. After extensive trials, Gaussian smoothing was chosen as the preferred method for generating target images, having demonstrated the most effective outcomes in terms of training efficiency and fidelity to the input data. The training of the model was conducted using the Adam optimizer with a learning rate set at 0.001 and a batch size of 16. This process was executed on a GeForce RTX 3090 graphics card.

### The creation of simulated datasets

The data collected comprises both simulated and real experimental datasets, encompassing seven categories in total. Simulated data includes sim-fish and DeepSpot datasets. The sim-fish data is generated using the Sim-FISH python package, which is part of the FISH-Quant v2[12] framework. The function `simfish.simulate_images()` is utilized for simulation, with different values set for each parameter to vary the signal noise and the number of signals in the images. DeepSpot dataset is created using a simulation script provided by the DeepSpot[22], with adjustments made to the parameters. All simulated data consist of images of size 512×512. Detailed parameter information can be found at this link: https://github.com/UFISH-Team/U-FISH/tree/main/data/simulation.

### The source of real experimental datasets

The real experimental data encompasses five categories: seqFISH, ExSeq, MERFISH, smFISH, and RCA. The seqFISH, ExSeq and MERFISH datasets are sourced from publicly available data at the following URLs, respectively: https://download.brainimagelibrary.org/biccn/zhangl/seqfish/Spinal_Cord_seqFISH_%2301-0001/, https://github.com/dgoodwin208/ExSeqProcessing/wiki/Example-Data-Set, and https://download.brainimagelibrary.org/02/26/02265ddb0dae51de/. The smFISH data is obtained from the deepBlink’s smfish.npz, available at https://figshare.com/articles/dataset/Datasets/12958037. The MiP-Seq data originates from the MiP-Seq[8]. All original images were cropped to a uniform size of 512×512, after which images of low quality, excessive noise or overexposed signals were removed to yield the final dataset required for this study.

### The measurement used in benchmark

To assess the quality of our results, a critical initial step is the matching of predicted signal points with the ground truth. To expedite this process, we construct a KD-Tree using the coordinates of the predicted points and then query this tree with the ground truth coordinates. This approach efficiently identifies the nearest predicted point for each ground truth signal. If the Euclidean distance between a matched pair (ground truth and predicted point) is less than a predefined cutoff (default value set at 3 pixels), the predicted point is classified as a True Positive (TP).

Once the number of TPs is determined, we calculate the False Positives (FP) by subtracting the number of TP from the total number of predicted points. Similarly, False Negatives (FN) are computed by subtracting the number of TPs from the total number of ground truth points. With these values: TP, FP, and FN, we can compute the F1 Score: 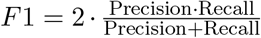 where 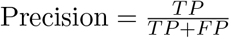 and 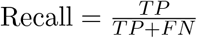.

In addition to the F1 Score, we also calculate the mean distance to evaluate the precision of localization. This metric is derived by averaging the distances between all matched pairs of points (ground truth and predicted). The mean distance serves as a measure of the localization error, offering an insight into the model’s performance in accurately pinpointing signal locations. This dual approach in metric calculation—focusing on both detection accuracy and localization precision—provides a comprehensive assessment of the model’s performance.

### Benchmark for deep learning methods

To evaluate the performance of U-FISH, we conducted a comparative analysis against three deep learning-based models: deepBlink[21], DetNet[19], and SpotLearn[20]. For deep-learning based approaches, due to the deep learning model’s inherent ability for transfer learning across diverse datasets, thus we employing a single model trained universally on all datasets to capitalize on its generalization capabilities. Notably, due to the lack of available implementations that could be readily retrained on our datasets for DetNet and SpotLearn, we re-implemented these models using the PyTorch framework. This re-implementation allowed us to integrate DetNet and SpotLearn seamlessly into U-FISH’s training and inference framework for a thorough testing. To ensure a fair comparison, we performed a grid search for DetNet’s hyperparameter *α* (the sigmoid shift parameter), aiming to identify the optimal setting for the validation dataset. This systematic approach allowed us to tailor DetNet’s performance closely to the specific needs of our evaluation, ensuring that all models were assessed under equitable conditions. As SpotLearn only predicts binary masks without actual point localization, the most efficient method for converting the characteristics of SpotLearn’s predicted results into point coordinates is by extracting the centroids of connected components. For DetNet, the predicted results consist of single pixels, and thus, the conversion solely involves mapping these single pixels to coordinates. We applied the `skimage.filters.threshold_otsu` function to binarize the enhanced output results generated by DetNet and SpotLearn networks. Subsequently, we employed the `skimage.measure.label` and `skimage.measure.regionprops` functions to identify signal points within the outputs. To ensure equitable comparisons, particularly considering the proclivity for dense signal overlap within SpotLearn’s target process, we integrated a watershed segmentation algorithm. More detailed parameters and training scripts can be found here: https://github.com/UFISH-Team/U-FISH/tree/main/benchmark#deep-learning-methods

### Benchmark for rule-based methods

U-FISH’s performance was benchmarked against ruled-based methods for single-molecule spot detection in images, including Big-FISH, RS-FISH[13], Starfish[29], and TrackMate[10]. To ensure a fair evaluation, we accounted for the limitation of rule-based methods, which cannot use a single set of parameters to achieve optimal performance across all datasets (Extended Data Fig.2c,d). Acknowledging this, we adopted a grid search approach to pinpoint the optimal parameters for testing. The evaluation metrics used for comparison were the F1-score and mean distance. Detailed grid search parameters and scripts can be found here: https://github.com/UFISH-Team/U-FISH/tree/main/benchmark#rule-based-methods

### Execution Time Measurement

Each of the methods was applied to a test dataset consisting of 550 images, each sized at 512×512 pixels. Execution time included tasks such as loading dependencies, loading weights, running the inference process, and logging information. This allowed us to estimate the average time needed for processing a single image. All time measurements were conducted on a workstation equipped with a GPU (NVIDIA GeForce RTX 3090) or a CPU (Intel Xeon 6330). To ensure reliability, each task was repeated five times on both the GPU and CPU to mitigate potential anomalies.

## Data availability

The U-FISH dataset is publicly accessible on HuggingFace at https://huggingface.co/datasets/GangCaoLab/FISH_spots. This release aims to support and encourage further advancements in FISH analysis research.

## Code availability

All code related to U-FISH has been made publicly available on GitHub. The U-FISH Python package can be found at https://github.com/UFISH-Team/U-FISH, offering the core functionalities and algorithms. For a graphical user interface experience, the U-FISH Web application is accessible at https://github.com/UFISH-Team/UFISH-Team.github.io, and the U-FISH Napari Plugin is available at https://github.com/UFISH-Team/napari-ufish, facilitating integration with the Napari viewer. Additionally, an online instance of U-FISH Web is hosted at https://ufish-team.github.io/, where users can directly engage in model predictions and dataset exploration without the need for local installation.

---

**Extended Data Fig. 1:**
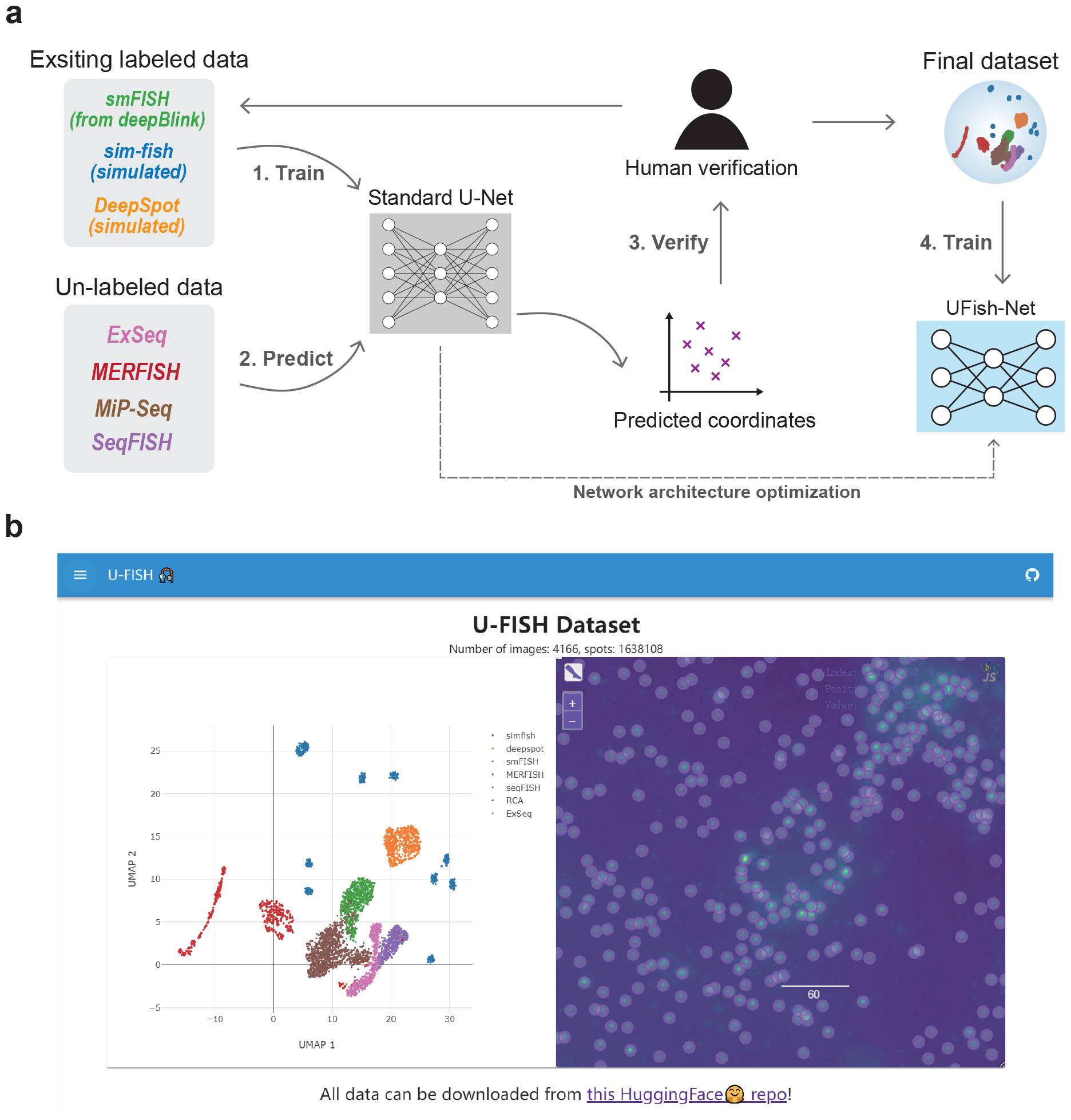
Dataset construction and browsing interface of U-FISH. **a**, This panel illustrates the process of constructing the U-FISH dataset. Initially, an auxiliary network with a standard U-Net architecture is trained using existing annotated datasets, including two simulated datasets and one smFISH dataset from deepBlink. This network is then utilized to pre-annotate an unlabelled dataset. Subsequently, human annotators meticulously review and correct these preliminary annotations, creating a new, fully annotated dataset. This dataset is then incorporated into the training set for retraining the network, and this iterative process continues until all datasets are annotated. In the final step, to enhance predictive performance, the U-Net architecture is refined and optimized to develop the UFish-Net architecture, which is then trained with the complete dataset. **b**, The U-FISH Web interface for dataset browsing and exploration is displayed. The interface is split into two sections. On the left, an interactive scatter plot created using Plotly.js shows the dimensionality reduction results of the U-FISH dataset images. When users click on a specific dot in the scatter plot, the corresponding image and its annotations are displayed on the right side in the Kaibu viewer. This setup provides an intuitive and user-friendly interface for users to navigate and examine the U-FISH dataset.

**Extended Data Fig. 2:**
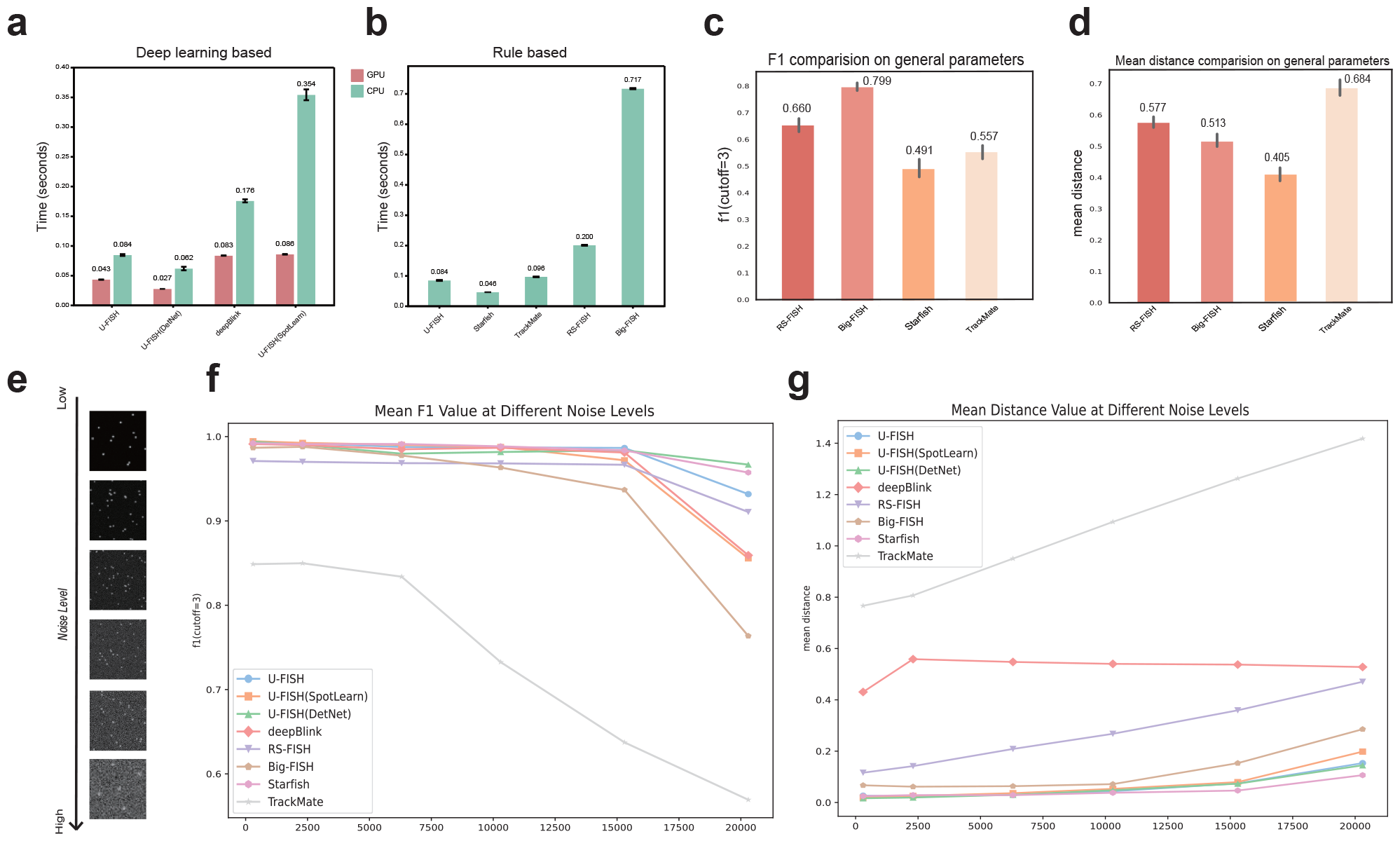
Benchmarks of spot detection methods: computational efficiency and accuracy across different conditions. **a**,**b**: The computational efficiency of spot detection methods. The y-axis represents the time required to process a 512 × 512 pixel test image. **a**, The results for deep learning-based methods. **b**, The result for rule-based methods. The color differentiation indicating CPU and GPU usage, the CPU tests were conducted on a node with two Intel Xeon 6330 (2.0GHz) processors, and the GPU tests on an NVIDIA RTX3090 with 24G VRAM. **c**,**d**, The performance of rule-based methods using a uniform set of optimal parameters across all datasets, focusing on mean F1 score (cutoff=3) and mean distance, respectively. These figures demonstrate the challenge for rule-based methods to adapt a single parameter set to various datasets effectively. **e**,**f**,**g**, assess the noise resilience of the methods. **e** displays a gradient of simulated noise. **f**,**g** then show each method’s performance under these noise gradients, evaluated using mean F1 score (cutoff=3) and mean distance metrics, respectively. This analysis highlights the differences in robustness and adaptability of the spot detection methods in response to varying noise levels.

**Extended Data Fig. 3:**
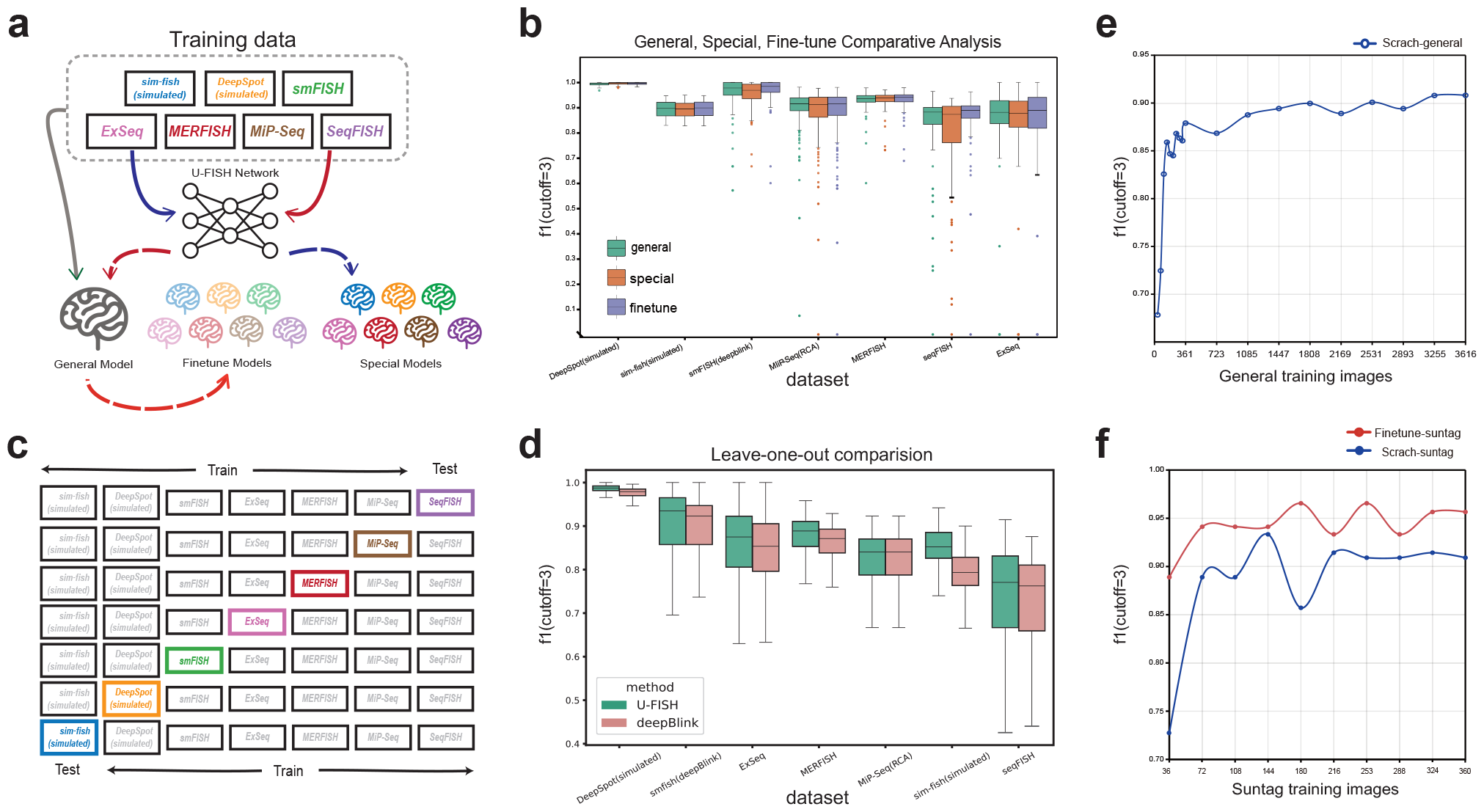
Model generalization and data requirement for fine-tuning.**a**, A schematic representation delineates the differences between General model, Finetune model, and Special model. The General model is trained uniformly using training data from all sources, while the Finetune model is further trained on top of the General model using data from specific sources. The Special model is trained from scratch with data exclusively from a single source.**b**, Performance comparisons of these three models across various training data sources. The Finetune model exhibits the best performance, followed by the General model, with the Special model lagging behind.**c**, An illustrative diagram explains the approach to test model generalization through a leave-one-out experiment. In this setup, data from one source is used as a test dataset while the remaining data sources form the training dataset. This tests the model’s performance on unseen images from the test dataset.**d**, Results of U-FISH and deepBlink methods in the leave-one-out test across different data sources are presented. This comparison highlights the generalization capabilities of each method.**e**, The impact of training data volume on model performance is depicted. The x-axis represents the number of 512×512 pixel training images, while the y-axis shows the resulting mean F1 score of the model. A notable improvement in performance is observed around 360 images, with the F1 score exceeding 0.9 when the number of images reaches over 1800.**f**, A comparison between training a model from scratch and fine-tuning based on the general model is illustrated. The x-axis shows the number of training images from Suntag data used, and the y-axis represents the achieved median F1 score. This graph demonstrates that fine-tuning on a general model with a smaller dataset can yield significant improvements in performance.

**Extended Data Fig. 4:**
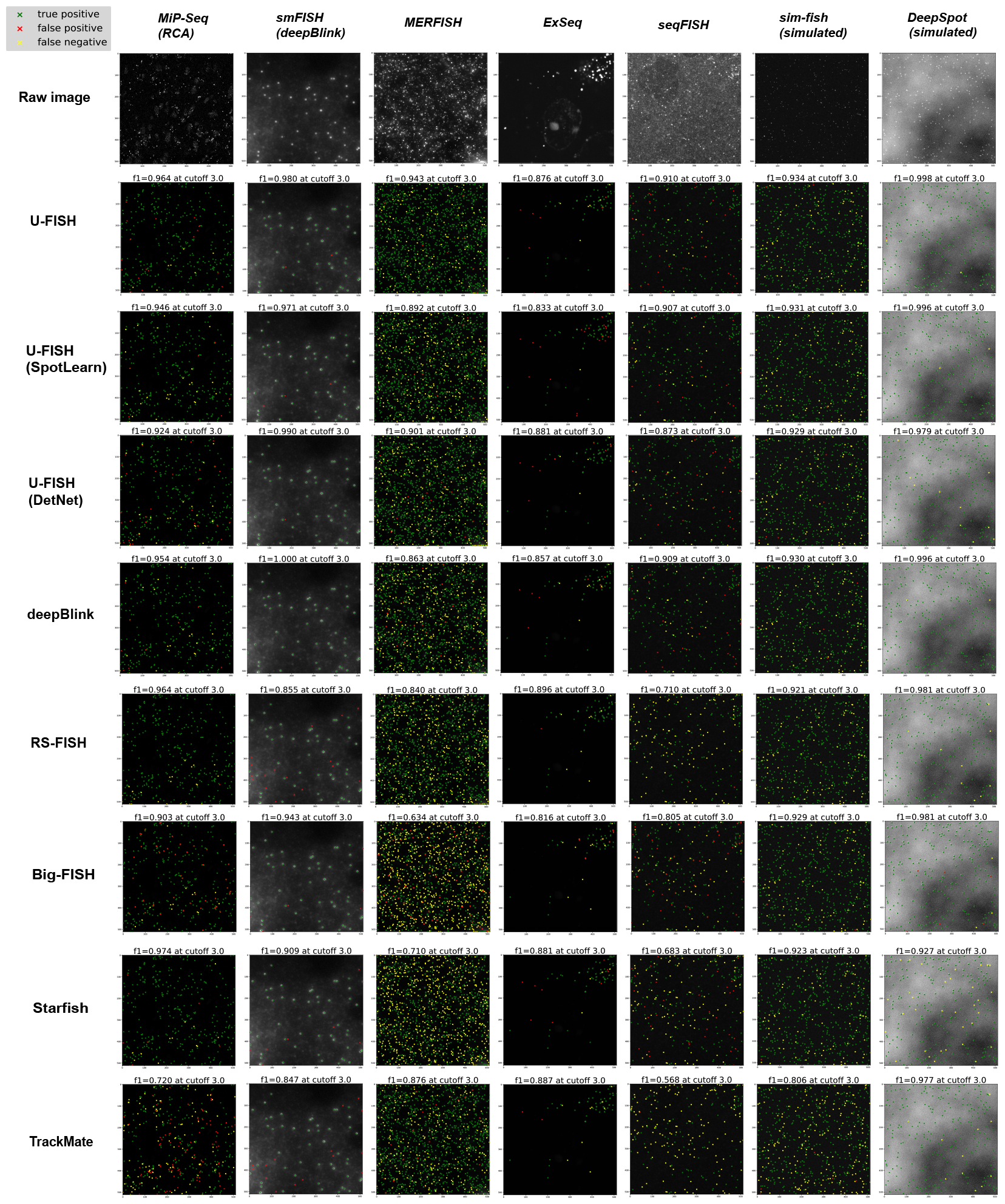
Benchmark examples with evaluation. This figure presents a series of example results from the benchmark tests, each representing different model and data type combinations. Within each example, the spots are annotated to distinguish between True Positives (TP), False Positives (FP), False Negatives (FN).

**Extended Data Fig. 5:**
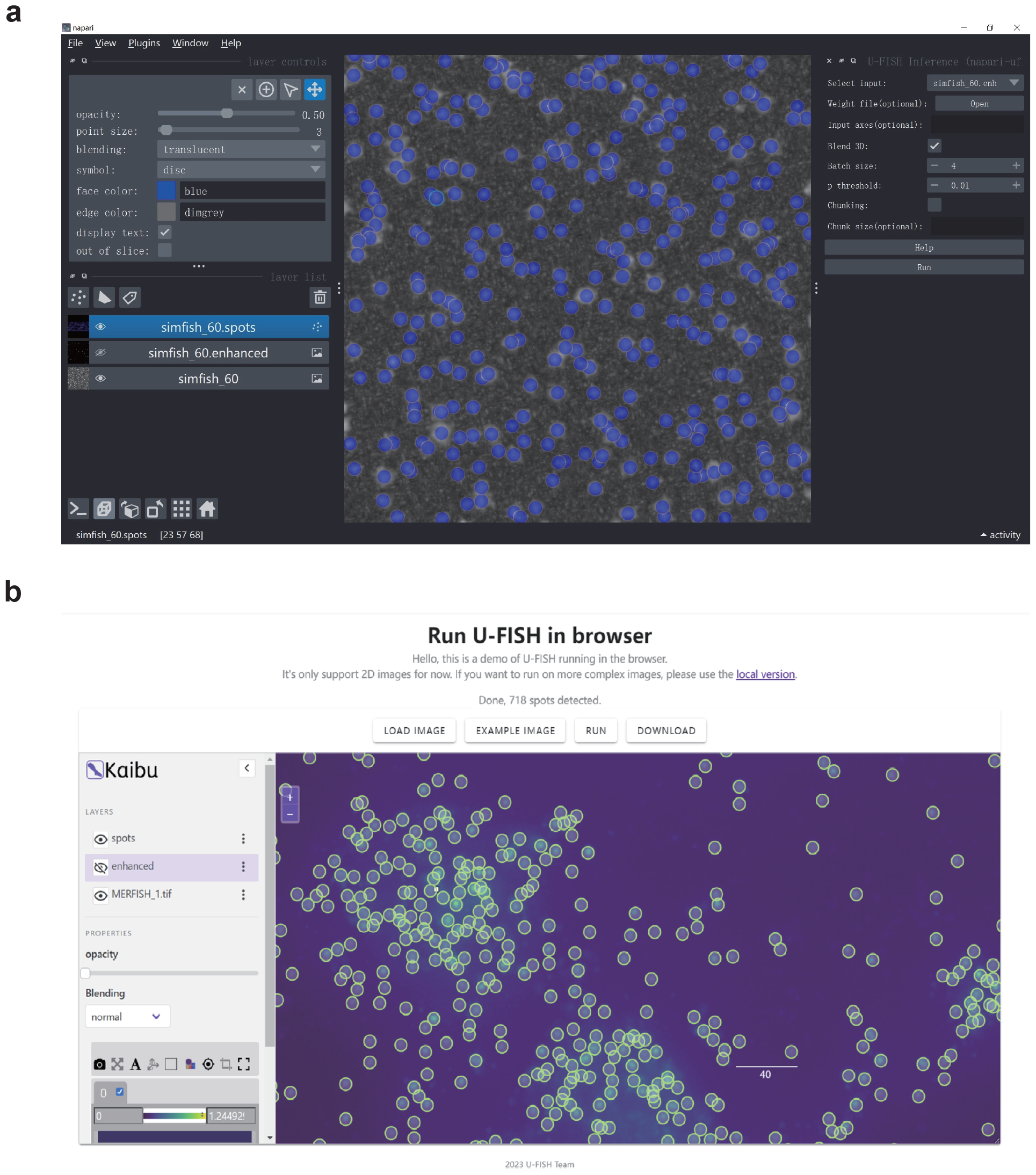
Graphical User Interfaces. **a**, The napari-ufish plugin features an Inference functionality interface, which, upon activation, displays a panel on the right side of the Napari viewer. This panel allows users to configure various operational options, such as enabling chunking and setting the chunk size. Upon clicking the “Run” button, the plugin employs the U-FISH model to compute and display the enhanced image and corresponding spots within the Napari browser, as illustrated. **b**, The web inference interface of U-FISH showcases an implementation using onnxruntime-web to run the U-FISH model directly in the browser. The interface integrates ImJoy[25] and Kaibu viewer for result visualization. It provides a straightforward, interactive user interface for uploading data, initiating the model run, and downloading the results. This web-based solution facilitates easy access and usage of U-FISH, allowing users to leverage its capabilities without the need for extensive setup or specialized software.

